# Morphological description of the male genital organs of *Pontoporia blainvillei* (Gervais & d’Orbigny, 1844)

**DOI:** 10.1101/2020.02.03.932897

**Authors:** Caroline Bizarre Randi, Lucas Guedes Spinelli, Renata de Britto Mari, Daniel de Souza Ramos Angrimani, Carolina Pacheco Bertozzi, Juliana Plácido Guimarães

## Abstract

*Pontoporia blainvillei* or Franciscana is a small cetacean endemic of South Atlantic Ocean and its highlighted as the most endangered species. Despite the necessity to development conservation strategies for *Pontoporia blainvillei* studies aiming to feature the reproductive biology and morphological characteristics of the genital organs are still scarce. In this context, the aim of this study was describe the morphological aspects of the genital organs of *Pontoporia blainvillei*. For this purpose, six male specimens of *P. blainvillei* stranded on the southeast region of Brazil, were used. Initially animals were measured, and the genital organs were collected, dissected and photodocumented. Then, fragments of all collected organs were submitted to microscopic analysis. The male genital organs of *P. blainvillei* showed similar results described for other cetaceans, such as, topography in abdominal cavity, presence of two testicles with epididymis, fibroelastic penis with sigmoid shape and presence of deferent duct. Moreover, a male uterus non-functional, two vestigial bones that support the penis, one retractor penis muscle and presence of associated muscles that support the entire reproductive apparatus were observed. However, some differences were present, such as, a simple cubic epithelium in the light of the epididymis, a small testicle even in sexually mature animals and absence of prostate, not previously observed in other cetaceans. Therefore, the animals analyzed showed similar features to other cetaceans, however some peculiarities were observed.

## Introduction

*Pontoporia blainvillei* (Gervais & d’Orbigny, 1844) is a small cetacean popularly known as Franciscana, is a Odontoceto from Pontoporiidae family, been endemic in South Atlantic Ocean [1]. Nowadays, the specie suffers an expressive population reduction due to industrial fishing and strandings [2, 3]. These factors configure the *P. blainvillei* as the most cetacean in risk in South America [3-5], which highlight the necessity to perform a conservation strategy, such as, the improvement of information about the reproductive biology of these animals.

Despite the importance of increase reproductive strategies for *P. blainvillei*, until this moment there are scarce information about the reproduction features of the specie. Franciscana is the one of the shortest life cycles among cetaceans, and although the sexual maturity are reached with two years old, the sexual dimorphism are exhibits only with four years old [6, 7]. Moreover, females had one gestation every one or two years, with a pregnancy period between 11 and 12 months [8]. Most females deliver birth on spring or summer, however some populations not present a defined season for reproduction [9-12].

In this context, new studies are required, because can provide several information to understand the Franciscana population dynamics. The definition of the reproductive features of *P. blainvillei*, for example, may allow the comprehension of modifications in testicles size during reproductive period, mating strategies and seasonal interactions, as described previously in other marine animals [2, 13]. This advance may provide the development of conservation strategies.

Therefore, the aim of this study was describe the morphological features of the male genital organs of *P. blainvillei*, using techniques of macroscopy and light microscopy, in order to contribute to the knowledge of reproductive biology of this specie.

## Materials and methods

### Animals

Six male specimens of *Pontoporia blainvillei* were used. The animals were provided by the Non-Governmental Organization, Project BioPesca. All of them were stranded and dead on beaches located on the coast of São Paulo, southeast region of Brazil. At macroscopically exam, the six animals were considered in normal condition despite the time of death.

### Sample Collection

Biometry of the animals was performed following the recommendations of the Protocol for the Conduction of Aquatic Mammals Strandings of the Northeastern Aquatic Mammal Strand Network - REMANE [14] to regulate the data collection for aquatic mammals. It was used a pachymeter to performed the testicular measures (width and length) from right and left testis.

The male genital organs was remove from abdominal cavity and then fixed in 10% formaldehyde solution, dissected and photodocumented for better analysis and description of results.

### Microscopic Analysis

For light microscopy analysis, the male genital organs, after fixation in 10% of formaldehyde solution, were dehydrated in increasing series of alcohols (70% to 100%), diaphanized in xylol and inserted in paraffin. Subsequently, samples were sectioned (6μm) and stained using the Hematoxylin-Eosin technique, for general visualization of the tissue. To determine the sexual maturity of *P. blainvillei* was analyzed the features of the seminiferous tubules, quantity of intertubular tissue and germinative epithelium.

### Statistical Analysis

Data from macroscopically and microscopically examination were descriptive. However, for the means and standard error of testicle (right and left) measurements (length and width) were used the SAS for Windows (SAS Institute Inc., Cary, NC, USA). The results are reported as untransformed means ± SEM.

## Results

The male genital organs of *P. blainvillei* were located in abdominal cavity and consist of a pair of testicles (located caudally to kidneys – Figure 1), epididymides, ducts deferens, urethra, male uterus and penis (Figure 1). The testicles (Figure 1 and 2) were elongated, in cylindrical shape, being that the mature animals showed the right testicle with a length of 4.2±0.3 cm and width of 1.4±0.4 cm, while the left testicle of the animals showed 3.9±0.6 cm in length and 1.2±0.5 cm in width (Table 1).

**Table 1.**
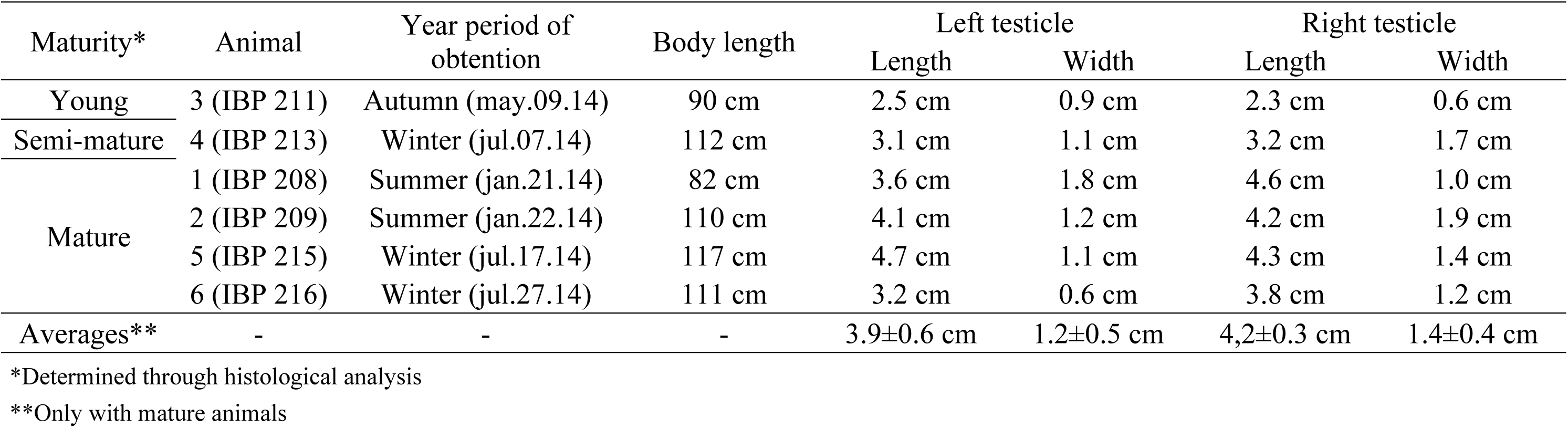
Correlation between the testicles (right and left) measurements (length and width) and the maturity of the animal, according to Hohn et al. [36].

**Fig 1.**
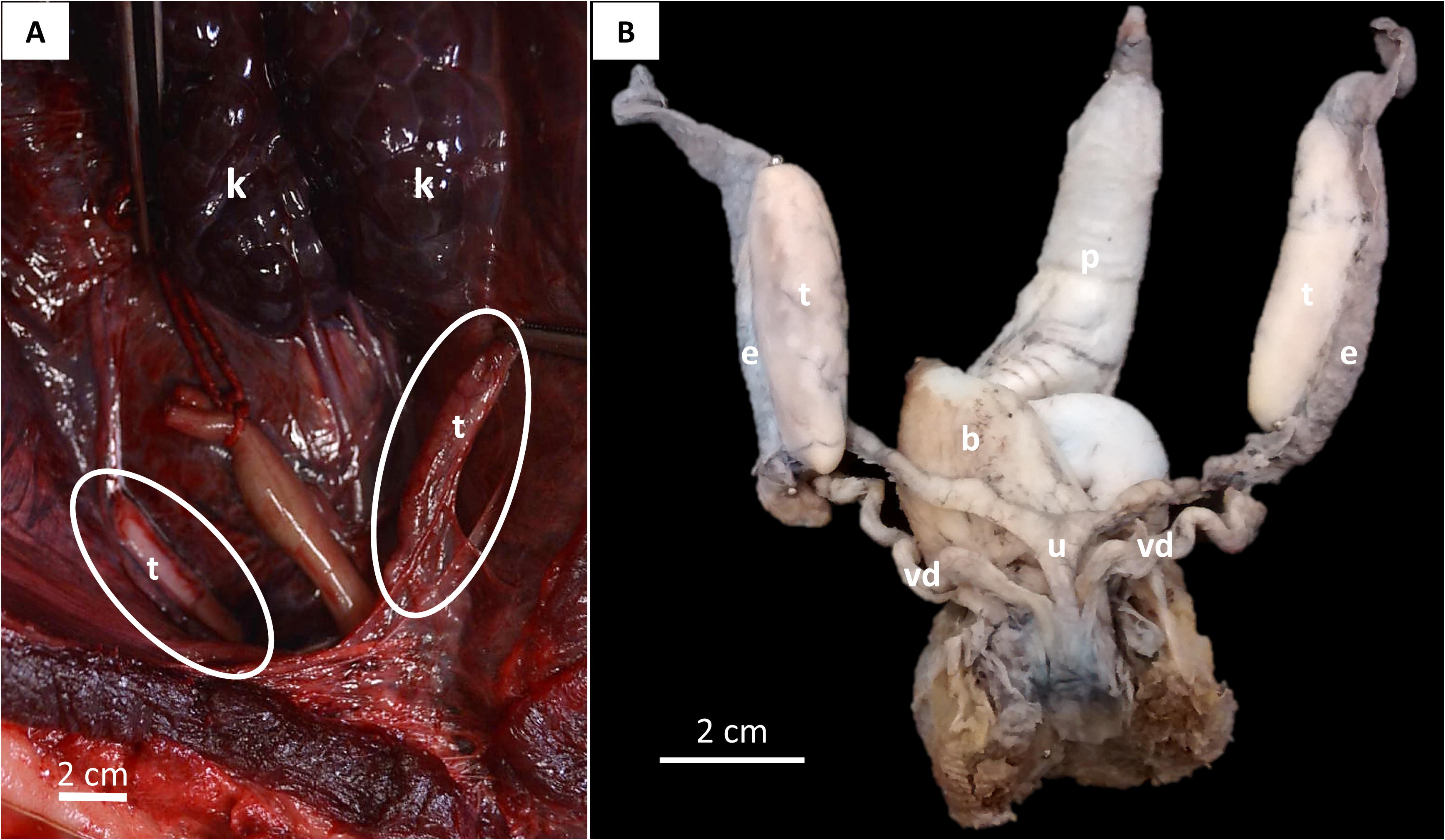
*Pontoporia blainvillei* male genital organs. A - Abdominal cavity of *P. blainvillei*, in their topography. B - Side view of testis and epididymis. Legend: t - testicle, R – kidney, e – epididymis, p – penis, u - male uterus, dd - vas deferens, b - bladder.

**Fig 2.**
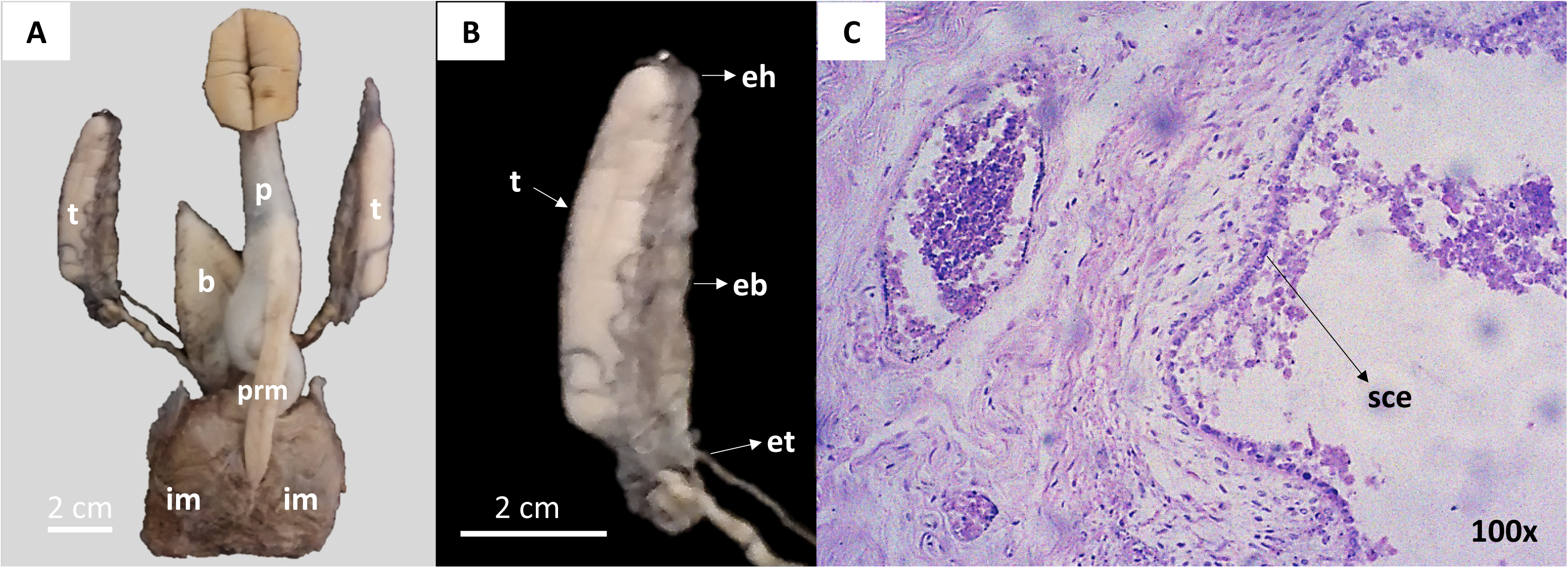
*Pontoporia blainvillei* male genital organs. A - Male genital organs of *P. blainvillei*. B – Testis and epididymis at higher magnification. C - Photomicrography of the *P. blainvillei* epididymis, which was possible to observe the simple cubic epithelium (ecs) in the tubular lumen. Legend: t – testis, cb – epididymis head, ca - epididymis tail, co – epididymis body, mrp - muscle retractor of the penis, mic - ischeocavernous muscle, b – bladder, p – penis, ecs - simple cubic epithelium.

Testicles were attached to abdominal cavity by mesorchium in lateral insertion. On the lateral side of each testis was located the epididymis (Figure 1 and 2). Epididymides were histologically characterized by a simple cubic epithelium in the tubular lumen. This organ was divided into head (cranial portion), body and tail (Figure 2).

Histologically, in the testis analysis, it was observed that immature animals had a great amount of connective tissue (intertubular tissue) in seminiferous tubules, absence of tubular light and presence of spermatogonias. In pubertal individuals, there was less space occupied by intertubular tissue, spermatogonia and spermatocytes (Figure 3). Moreover, in mature males, a small amount of intertubular tissue was observed, resulting in reduced space between seminiferous tubules. The germinal epithelium showed all stages of maturation, including spermatids (Figure 3).

**Fig 3.**
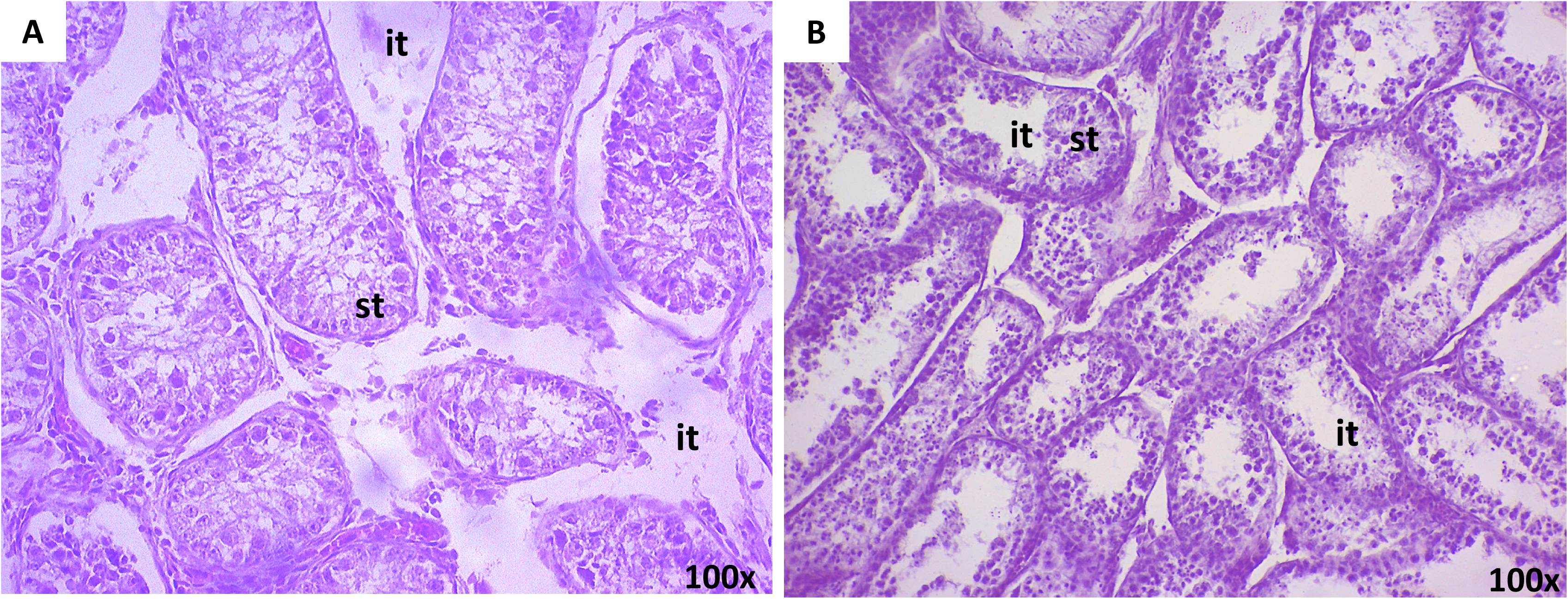
*Pontoporia blainvillei* male genital organs. A - Photomicrography of the testis of a pubertal individual of *Pontoporia blainvillei* by Hematoxylin-Eosin. B - Photomicrography of the testis of a mature individual of *Pontoporia blainvillei* by Hematoxylin-Eosin. Legend: ts - seminiferous tubule, si - intertubular tissue (connective tissue).

Deferens ducts were retroperitoneal, leading caudally to urethra, and extremely convoluted in the region proximal to testis. Histologically, ducts deferens had a narrow light, consisting of muscle layers, smooth connective tissue, and a pseudostratified columnar epithelium. The presence of a male uterus located between the deferens ducts and dorsal to bladder (Figure 1) was observed, which, histologically, presented a simple cubic epithelium (Figure 4).

**Fig 4.**
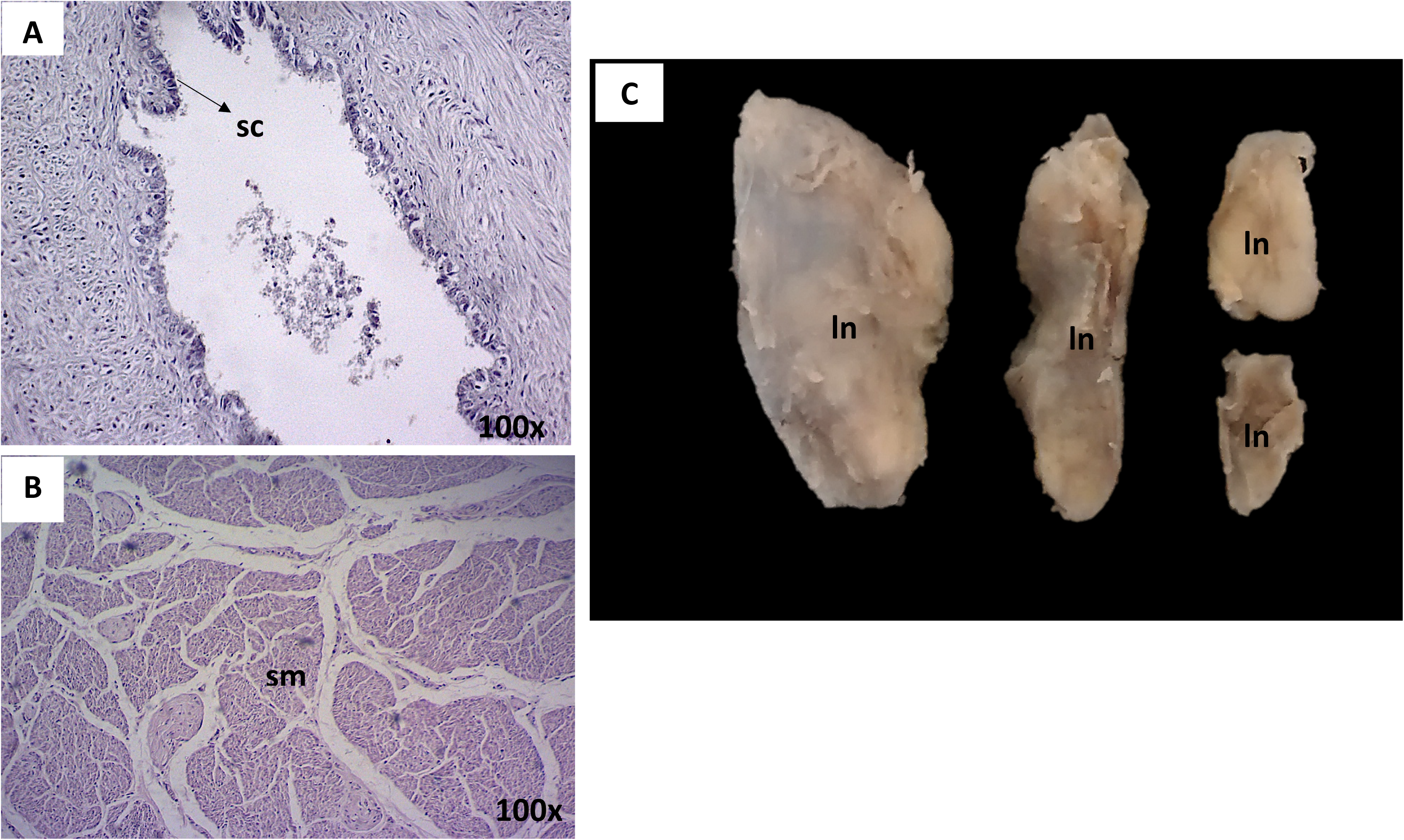
Pontoporia blainvillei male genital organs. A - Photomicrograph of the male uterus of Pontoporia blainvillei by Hematoxylin-Eosin. B - Photomicrography of the retractor muscle of the penis by Hematoxylin-Eosin. C – Prostate lymph nodes. D - Photomicrography of a Pontoporia blainvillei lymph node by Hematoxylin-Eosin. Legend: ecs - simple cubic epithelium, fml - smooth muscle fibers, lin – lymph nodes.

The *P. blainvillei* had a fibroelastic penis (Figure 5) which in quiescent state presented a sigmoid shape. It is covered by prepuce and had the retractor muscle of the penis, consisting of smooth muscle fibers (Figure 2 and 5). Histologically, penis had a albuginea tunic with a high resistance composed by connective tissue with large amount of collagen. Penis was formed by cavernous body, and it is possible to observe a transitional epithelium in its tubular light referring to urethra. Prepuce is formed by a retractile fold of penis skin containing connective tissue, and pavement-stratified epithelium (Figure 5).

**Fig 5.**
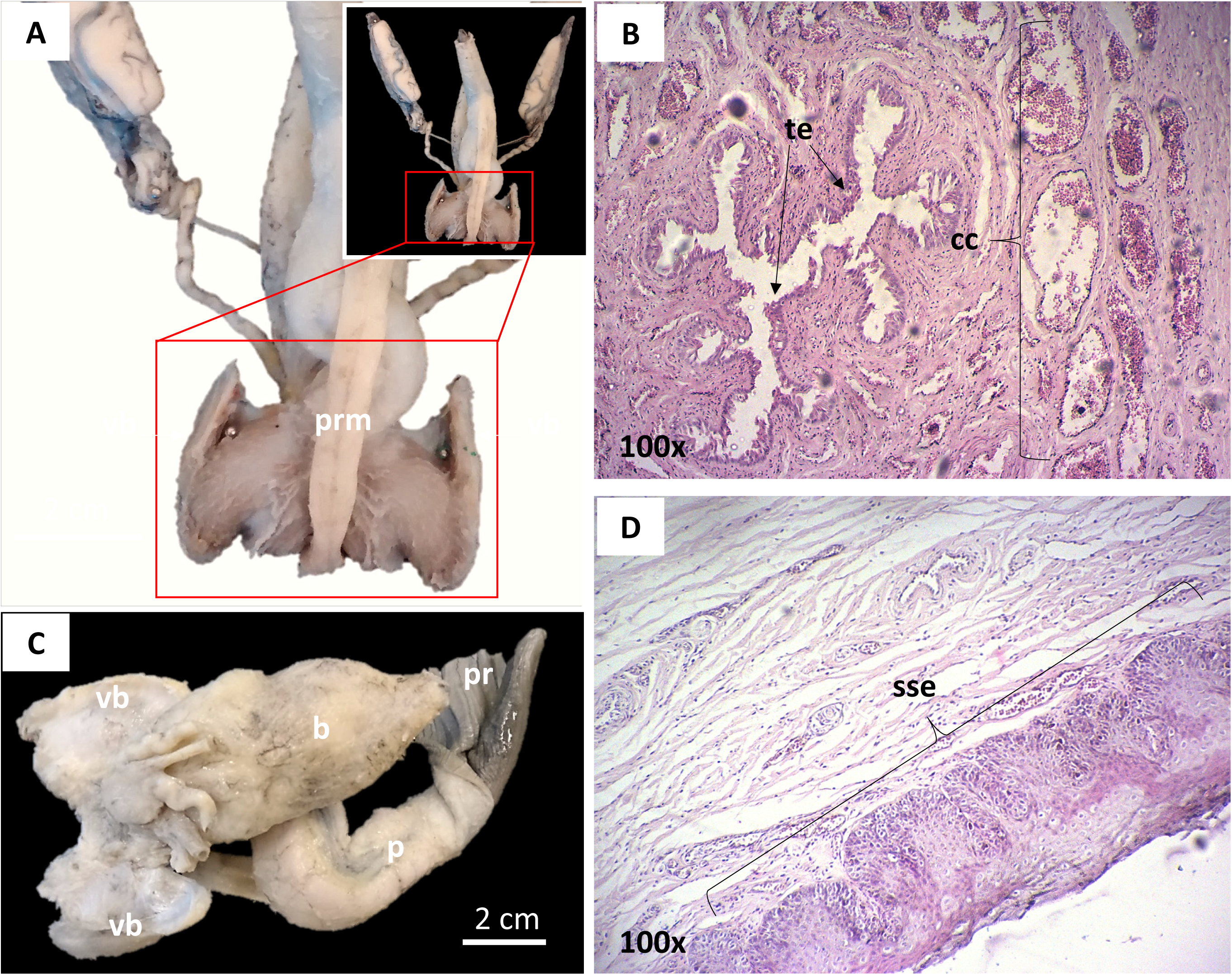
*Pontoporia blainvillei* male genital organs. A - Male genital organs of *P. blainvillei*. B - Photomicrograph of *Pontoporia blainvillei* penis by Hematoxylin-Eosin. C - Male genital organs of *P. blainvillei* demonstrating the penis prepuce. D - Photomicrography of *P. blainvillei* prepuce by Hematoxylin-Eosin. Legend: t – testicle, cb – epididymis head, ca - epididymis tail, co – epididymis body, mrp – penis retracted muscle, o - vestigial bones, p – penis, b – bladder, pr – prepuce, etr - transitional epithelium, cc - corpus cavernosum, eep - stratified squamous epithelium.

Male reproductive tract was supported by a pair of vestigial bones (Figure 5), which were covered by ischiocavernous muscle, featured by striated muscle fibers. These vestigial bones were remnants of pelvic bones in which the penile pillars were inserted. Lymph nodes were located into the prostate compressor muscle (Figure 4), however the prostate was not detected.

## Discussion

In this study, were evaluated the morphological features of the male genital organs of *P. blainvillei* (Gervais & d’Orbigny, 1844) by macroscopic and microscopic analysis in order to contribute to the knowledge of reproductive biology of this specie.

Despite some peculiarities, the male reproductive tract of *P. blainvillei* had a greater similarity to other cetaceans, as the topography in the abdominal cavity [15]. In the most mammals, testicles develop occurred inside abdominal cavity and migrate through inguinal canals to scrotum [16]. On the other hand, testicles from *P. blainvillei*, as the other cetaceans, are located on abdominal cavity, which promotes advantage to the animal hydrodynamics and suggests a wide importance for body thermoregulation [17].

Testicles of *P. blainvillei* were in pairs, with cylindrical and sharp poles as described previously in other cetaceans [17, 18]. In Odontoceti, testicles reach a considerable size in sexually active adults [19]. However, in Pontoporiidae testicles were characterized by small size, even in mature individuals, which may be related to the reproductive strategy of the species. In *P. blainvillei*, testicles (left and right) presented symmetry, as previously observed for the specie [20].

Testis of cetaceans are connected by mesorchium in abdominal cavity, in *P. blainvillei* was observed a lateral insertion, similar to *Stenella sp.* [21] and *Tursiops truncatus* [22]. Histologically, testis structure was similar to described in cetaceans [17] and the seminiferous tubules observed were typical to mammals [23]. In this study, all animals analyzed, despite the age, presented the size and distance of seminiferous tubules, amount of intertubular tissue and presence of sperm cells followed the pattern already described [24].

On the lateral side of each testicle, the presence of epididymis were observed, such as, the presence of head, body and tail, observed previously in cetaceans [15, 25, 26] and in domestic mammals [27, 28]. Moreover, epididymis of *P. blainvillei* are characterized to be fixed in testis, as in *Phocoena phocoena* [22]. Histologically, epididymis of *P. blainvillei* presented a single convoluted duct and a simple cubic epithelium [17, 29].

Attached to epididymis final portion are located deferens ducts, which is responsible for sperm transportation and in *P. blainvillei* had the same pattern observed in cetaceans and mammals [29, 30]. The deferens ducts of *P. blainvillei* are extremely convoluted in proximal region of each testicle, and more rectilinear in distal region. Although this ducts deferens characteristic are present in the most cetaceans [31] it is different in *Cephalorhynchus commersonii* [32].

Microscopically, it was observed that deferens ducts presented a limit light and a thick wall, consisting of connective tissue, smooth muscle and pseudostratified columnar epithelium. This is the first description about the microscopically features of deferens ducts in *P. blainvillei.* In general, are few descriptions about the histology of vas deferens on cetaceans, and among them there is a controversy regarding the epithelial type. In *Pseudorca crassidens*, for example, are described as columnar epithelium with one or two layers [17] and in *Stenella frontalis* simple ciliated epithelium [21]. On the other hand, in mammals the deferens ducts is characterized as a pseudostratified [23].

In the most distal portion of deferens ducts in *P. blainvillei* it was observed a union between ducts and male uterus in dorsal region of the bladder. This fact also occurs in *Tursiops truncatus* [22] and *Stenella frontalis* [21]. Also, the male uterus is a remnant of Müller duct [23]. In the animals analyzed, male uterus is a small, tapered and tubular structure, as observed in *Pseudorca crassidens* [17]. However in other cetaceans it can be extensive and bicorn, as in *Mesoplodon sp.* [17] or even absent, as in *Tursiops truncatus* [22]. Histologically, male uterus presented a simple cubic epithelium. The only histological descriptions of male uterus in cetaceans are in *Pseudorca crassidens* [17], which was similar to our data in *P. blainvillei.*

The *P. blainvillei* penis is fibroelastic, remains in completely quiescence state inside the abdomen, being exposed only during erection, which makes it similar in location and structure to other cetaceans [29]. Moreover, two vestigial bones were observed in *P. blainvillei*, in which the penile pillars were inserted. These vestigial bones were inserted in a ischaocavernous muscle [17]. This feature is not only noted in cetaceans, but in all permanently aquatic mammals [31, 33].

In *Delphinus delphis* there is a substantial increase in the size of vestigial bones when animals reach sexual maturity [24]. In fact, our data showed a larger size in vestigial bones in adult individuals than in young, which may be related to the support the penis function. Furthermore, in addition to ischemic muscle, the *P. blainvillei* present the prostate muscle compressor, despite the prostate gland was not detected.

In our samples of *P. blainvillei* it was possible to observe that all animals presented sigmoid penis, commonly in animals with fibroelastic penis, with a spiral sigmoid flexure. Thus, function, structure and location are similar to other marine mammals [21]. Moreover, in some domestic mammals this muscle are present in pair [34], however in all *P. blainvillei* analyzed the muscle was unique. Histologically, the penis retractor muscle is featured by smooth muscle fibers, as described in *Stenella frontalis* [21] and *Pseudorca crassidens* [17].

The penis was pointed and elongated, no gland were observed as described in other cetaceans [17, 29]. However, other penis forms were reported, such as, sharp in *Physeter catodon* [35] and tongue format [21]. Microscopically, a transitional epithelium was observed in the penis corresponding to urethra, this histological pattern had already described for other cetaceans [31].

The prepuce of *P. blainvillei* basically consist of a retractile fold of penile skin containing dense connective tissue, and is composed of a paved epithelial tissue. Furthermore, in *P. blainvillei* the prepuce has a genital cleft in the abdomen where the penis is exposed when erectile, the same features were observed in other cetaceans [17].

## Conclusion

In conclusion, the description of these morphological features presented in our study can be used as a basis for research on the reproductive biology of *P. blainvillei*. This data may encourage future studies focusing on conservation strategies for this species. In addition, anatomical and functional knowledge of genital organs of this species allows the development of reproductive biotechnologies, which are extremely important for the conservation.

